# A hydraulic instability drives the cell death decision in the nematode germline

**DOI:** 10.1101/2020.05.30.125864

**Authors:** N. T. Chartier, A. Mukherjee, J. Pfanzelter, S. Fürthauer, B. T. Larson, A.W. Fritsch, M. Kreysing, F. Jülicher, S. W. Grill

## Abstract

Oocytes are large and resourceful. During oogenesis some germ cells grow, typically at the expense of others that undergo apoptosis. How germ cells are selected to live or die out of a homogeneous population remains unclear. Here we show that this cell fate decision in *C. elegans* is mechanical and related to tissue hydraulics. Germ cells become inflated when the pressure inside them is lower than in the common cytoplasmic pool. This condition triggers a hydraulic instability which amplifies volume differences and causes some germ cells to grow and others to shrink. Shrinking germ cells are extruded and die, as we demonstrate by reducing germ cell volumes via thermoviscous pumping. Together, this reveals a robust mechanism of mechanochemical cell fate decision making in the germline.

## Main Text

Oocytes are cells that both transmit genetic information to the next generation and provide sufficient cellular material to develop a fertilized zygote into an embryo (1). During their maturation from primordial stage future oocytes undergo extensive growth, which is often achieved at the expense of other germ cells that ultimately die by apoptosis (2,3). This altruistic behaviour and the associated exchange of material is facilitated by a syncytial structure with a shared cytoplasm. While apoptosis in the germline can serve a quality control function by removing damaged cells (4), the vast majority of germ cells that undergo apoptosis under normal conditions appear healthy (5). A fundamental question therefore is: out of a seemingly homogeneous population of healthy germ cells, how are cells selected to live or die?

The germline of the adult hermaphrodite nematode *Caenorhabditis elegans* (*C. elegans*) captures all the essential features to address this open question. It is a tubular syncytium consisting of germ cells that surround a central cytoplasmic compartment called rachis, to which all the germ cells are connected via openings called rachis bridges (Fig. 1A) (6,7). Germ cells originate in a mitotic zone from a pool of stem cells residing in the distal tip of each gonad arm, and undergo meiotic maturation as they move towards the proximal turn (6). During this progression some of the germ cells grow to become oocytes while the rest shrinks and dies by physiological apoptosis (Fig. 1A) (5). Although the core apoptotic machinery was shown to drive the final steps of cell death, the mechanisms that select and initiate apoptosis in individual germ cells are still unclear (4,8).

**Fig. 1:**
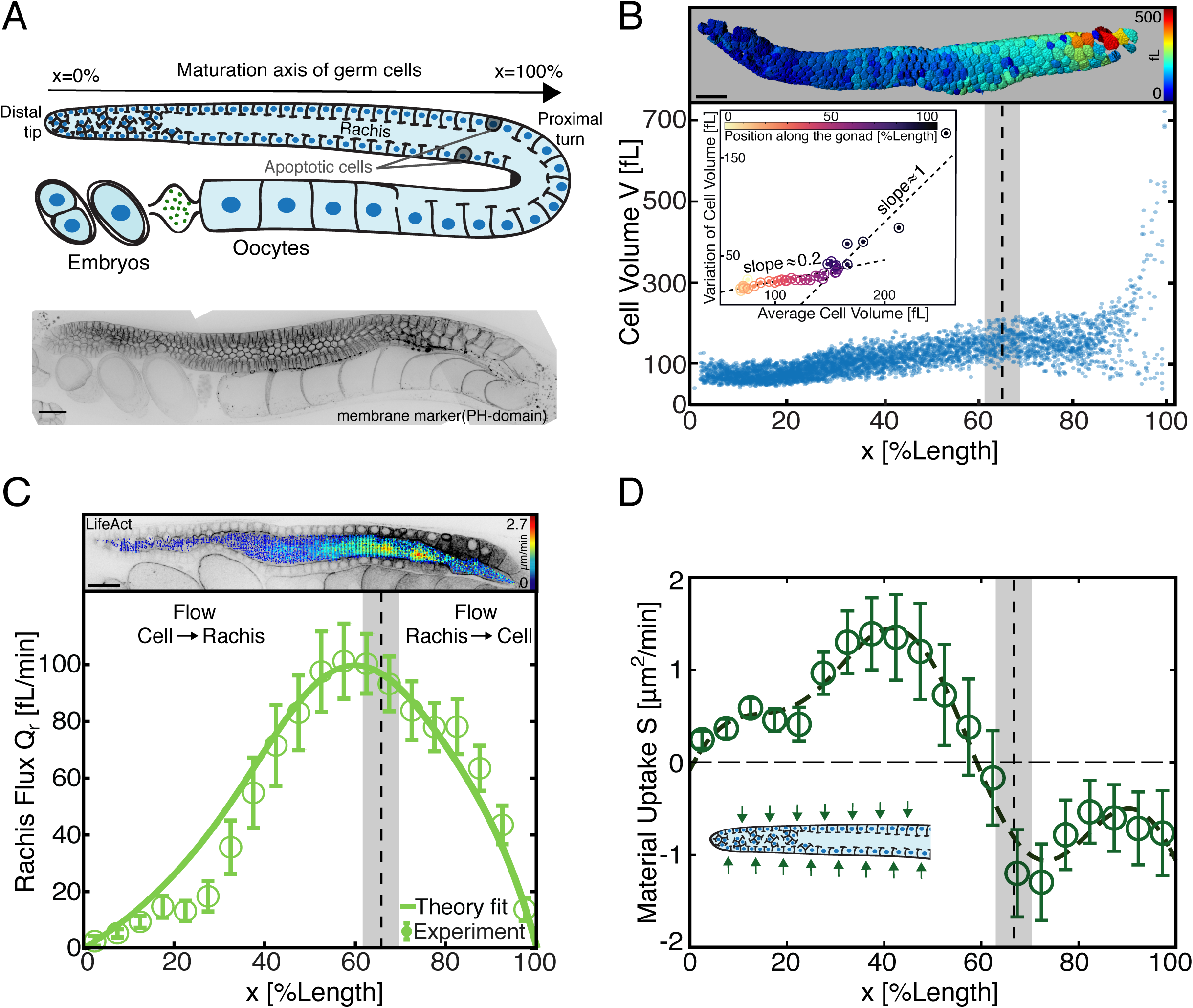
Volumes and fluxes in the *C*.*elegans* gonad. **A**, Top, schematic of an adult hermaphrodite gonad arm. Bottom, representative fluorescence image of a gonad expressing the membrane marker mCherry::PH(PLC1delta1). **B**, Top, germ cells color-coded according to cell volume. Bottom, germ cell volume V along the gonad from distal tip (0% length) to proximal turn (100% length), for 5265 germ cells from 18 gonad arms. Inset, standard deviation (SD) plotted against the average of germ cell volumes determined at 40 color-coded positions along the gonad. We observe two different relationships between SD and average, indicative of a transition between two growth modes. Dashed vertical lines and grey boxes in B-D denote the position and associated confidence interval where the distribution of germ cell volumes is no longer unimodal. **C**, Top, Cross-section of a gonad expressing Lifeact::mKate overlaid with color-coded flow speeds as obtained by PIV (see supplement). Bottom, cytoplasmic flux *Q*_*r*_ through the rachis as a function of position along the gonad. Open circles, *Q*_*r*_ determined from PIV speed distributions obtained from 10 gonad arms. Solid line, best parameter theory fit given the profile of material uptake *S* shown in D. **D**, Open circles denote material uptake *S* into the gonad from the outside, determined by volume conservation of rachis flux (C) and volume flux associated with germ cells moving from distal to proximal (Fig. S1E). Dashed line shows a smoothened representation of the material uptake profile (see supplement). Inset, green arrows indicate material uptake from the surrounding. Scale bars; 20 *µm*. Error bars indicate the error of the mean at 95% confidence.

Oocyte growth in *C. elegans* has been shown to rely on long range cytoplasmic streaming (9), but how and why germ cells shrink remains elusive. To identify a potential relationship between germ cell growth, shrinkage, and apoptosis, we first set out to both quantify where along the gonad germ cells grow, and characterize how this growth proceeds (Fig. 1B). Confocal imaging followed by 3D-membrane-based segmentation of adult germlines expressing the membrane marker mCherry::PH(PLC1delta1) allowed us to measure individual germ cell volumes along the distal to proximal axis until the turn region (Movie S1). We find that germ cells near the distal tip have a volume of approximately 100 fL. As germ cells mature along the gonad they first grow collectively in volume to approximately 150 fL (Fig. S1A). Prior to the proximal turn the variation of germ cell volumes increases drastically, and germ cells range from very small (∼ 65 fL) to very large sizes (∼ 1200 fL). While the distribution of germ cell volumes is unimodal in the distal region, it becomes bimodal close to the turn (Figs. 1B, S1B). This suggests a transition from a homogeneous to a heterogeneous growth mode of germ cells along the gonad. In order to identify the precise location where this transition occurs, we investigated the average and the standard deviation of germ cell volumes in different regions along the gonad (Fig. 1B, bottom inset). Both quantities appear to be linearly related but the associated slope changes sharply, which can be used to locate the transition zone to 65% *±* 3.75% germline length. Two alternative methods to identify the transition point gave a similar result (see supplement). Note that physiological apoptosis occurs proximal to this transition point, from about 70% to 90% germline length (Fig. S1C).

Both the homogeneous and the heterogeneous mode of germ cell growth must rely on the addition of cytosolic volume. Germ cells can either grow by receiving material from the rachis inside or from the surrounding tissue outside, such as the intestine (10). We set out to identify for the two regions of growth where the corresponding cytosolic volume comes from. For this, we determined the flux of cytoplasm through the rachis along the gonad. Due to incompressibility, an increase in rachis flux along the gonad implies that germ cells contribute material to the rachis, while a decrease means that germ cells receive material from the rachis. We performed Particle Imaging Velocimetry (PIV) (11) on mid-plane confocal sections of the germline expressing LifeAct::mKate to determine a cytoplasmic velocity field inside the rachis, which we then used to infer the rachis flux as a function of position along the germline (Fig. 1C, Movie S4, see supplement). We find that the flux of cytoplasm through the rachis increases monotonically along the distal part of the gonad, peaks at approximately 60% germline length, and decreases thereafter. Hence, germ cells donate material to the rachis prior to 60% germline length, while they receive material from the rachis thereafter.

Interestingly, germ cells grow prior to 60% germline length (Figs. 1B, S1A) despite losing cytoplasm to the rachis. This implies that they must be receiving material from the outside. We inferred the profile of material uptake (Fig. 1D) using volume conservation, considering the rachis flux and the volume flux associated with germ cells moving from distal to proximal (see supplement). We find that in the distal region and up to approximately 60% gonad length, material uptake is positive and germ cells grow by receiving material from the outside. Material uptake becomes negative beyond 60% indicating a loss of material to the outside, possibly via removal of apoptotic cells. We conclude that the material associated with homogeneous growth of germ cells comes from outside, while the heterogeneous growth mode is associated with germ cells receiving cytoplasm from the rachis.

Pressure differences between germ cells and rachis drive cytoplasmic exchange through rachis bridges. To discuss the hydraulics of the gonad, we construct a one-dimensional physical model that relates pressure profiles to flows of germ cells and rachis cytoplasm as well as material exchange between germ cells and rachis (Fig. 2A). Using the profile of material uptake (Fig. 1D), this theory recapitulates rachis and germ cell fluxes (indicated by solid lines in Figs. 1C, S1E) and predicts a profile of cell to rachis current that matches the experimental estimates (Fig. 2B). Consistent with the observation that the rachis flux peaks around 60% gonad length, this current changes sign at the same location (Fig. 2B). Because pressure differences drive cell to rachis currents, the pressure difference between cells and rachis *P*_*c*_ −*P*_*r*_ also changes sign at this location. We next asked if this inversion of the pressure difference might be at the heart of the transition from homogeneous to heterogeneous mode of germ cell growth. We note that the change of a monomodal to a bimodal volume distribution is indicative of an instability of germ cell volumes during growth. A similar instability arises when blowing simultaneously into two rubber balloons in the attempt to inflate them both. Here, only one balloon inflates. Because the larger balloon can be inflated at lower pressures than the smaller one, the situation where both simultaneously inflate is mechanically unstable (12).

**Fig. 2:**
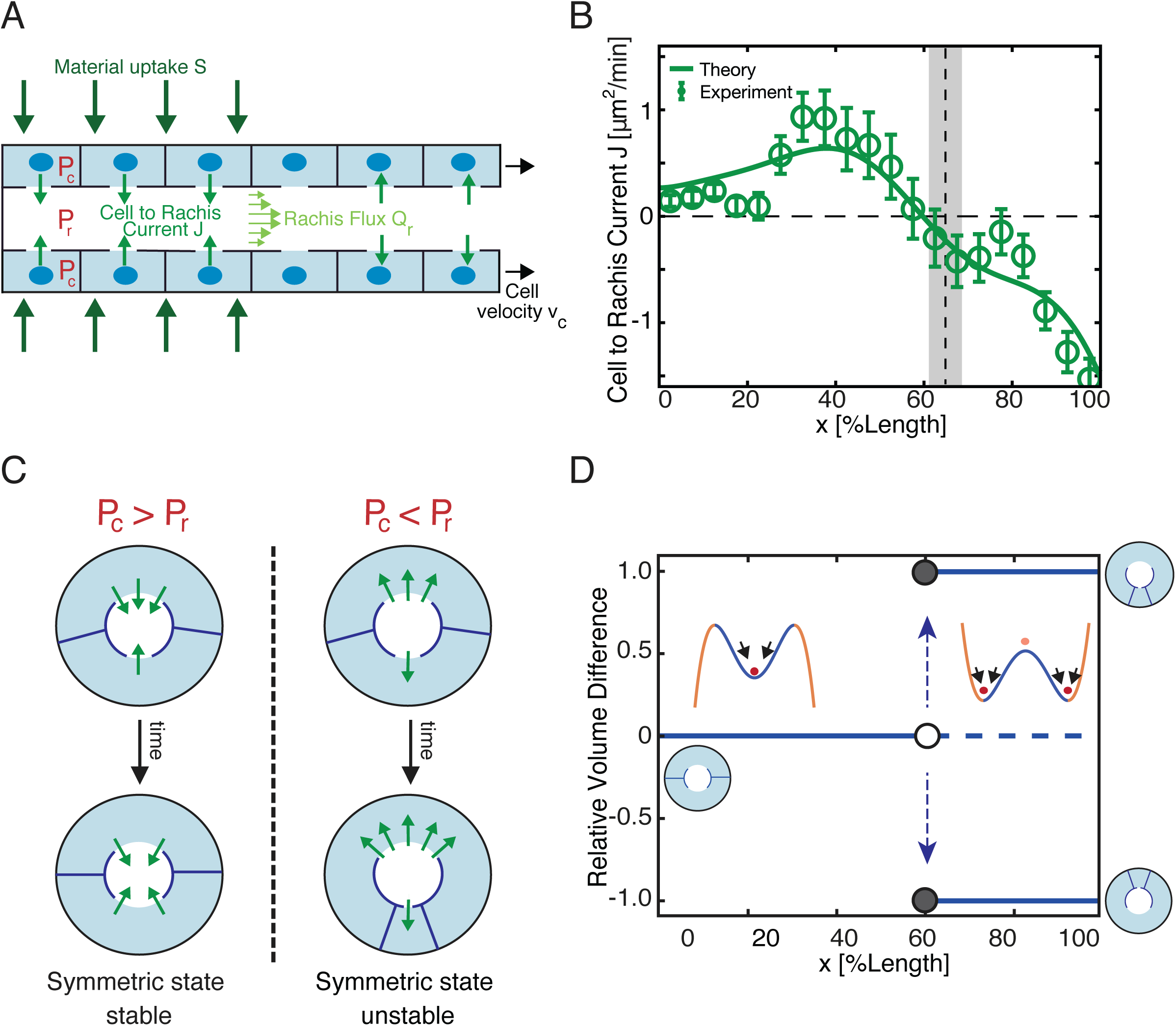
A theoretical model of germ cell and rachis fluxes reveals a hydraulic instability. **A**, Schematic of a 1D hydrodynamic model for pressures, material fluxes and volume exchange in the *C. elegans* gonad. *P*_*c*_ and *P*_*r*_ denote the pressure field in germ cells and rachis, *S* the profile of material uptake from the outside, *J* the germ cell to rachis current associated with flows through rachis bridges, *Q*_*r*_ the rachis flux and *v*_*c*_ the germ cell velocities. **B**, Green open circles, estimated cell to rachis current *J* along gonad length, vertical dashed line and grey bar denote the region of transition between germ cell growth modes (Fig. 1B). Solid line, best parameter theory fit given the profile of material uptake *S* shown in Fig. 1D. **C**, Evolution of small volume differences between coupled germ cells with time. The symmetric state of equal germ cell volumes is unstable when the pressure in the rachis is higher than in germ cells (right). Here, the initially larger cell grows at the expense of the smaller one. **D**, The cell volume difference in a cell doublet bifurcates around 60% germline length where the pressure difference changes sign. Insets illustrate cell configurations and corresponding pseudo-potentials (see supplement). Error bars indicate the error of the mean at 95% confidence.

Could such an instability also arise in the germline (13-18)? We consider a simplified configuration of two germ cells with slightly different volumes that surround the rachis (Fig. 2C). If the pressure is larger in the rachis than in the germ cells, we have a situation that is similar to the balloon scenario. To discuss the hydraulics of this simplified two-cell system, we take into account that large germ cells tend to have larger rachis bridges than small germ cells (Fig. S1F). A linear stability analysis of germ cell volumes reveals that the state of equal germ cell volumes is stable only if the pressure inside germ cells is larger than in the rachis (see supplement, Fig. 2C). An instability occurs when the pressure in the rachis is larger than in germ cells: a small difference in germ cell volumes will increase, leading to the growth of the larger germ cell at the expense of the smaller one. Importantly, this hydraulic instability is expected to be triggered when the pressure difference between rachis and germ cells becomes negative at 60% gonad length (Fig. 2D), beyond which it would drive a heterogeneous mode of germ cell growth as is observed in this regime (around 65% gonad length, see Fig. 1B).

This hydraulic instability presents a possible mechanism by which germ cells become fated to die. It generates a few large cells at the expense of smaller shrinking cells in a coarsening process. Apoptosis is then triggered in shrinking cells, which leads to their removal. However, an alternative scenario is that unknown molecular signals first induce apoptosis, which subsequently leads to the shrinkage of those cells fated to die. To test this alternative possibility, we inhibited apoptosis of germ cells by RNAi (19) targeted against the caspase *ced-3* (4,5) and evaluated if germ cells still shrink. We find that in the absence of apoptosis germ cells are no longer removed (5), however, we still observe that some cells in the proximal region shrink (blue dots in Fig. 3A). Furthermore, similar to unperturbed conditions, *ced-3(RNAi)* gonads show a transition from a homogeneous to a heterogenous mode of growth (grey bar in Fig. 3A), and the position of this transition is close but a bit proximal to the location where both the cell to rachis current and the pressure difference between germ cells and rachis changes sign (Fig. 3A), and where the rachis flux peaks (Fig. 3B). Together, this eliminates apoptosis as the cause of germ cell shrinkage and supports the idea that germ cell fate is determined by a hydraulic instability.

**Fig. 3:**
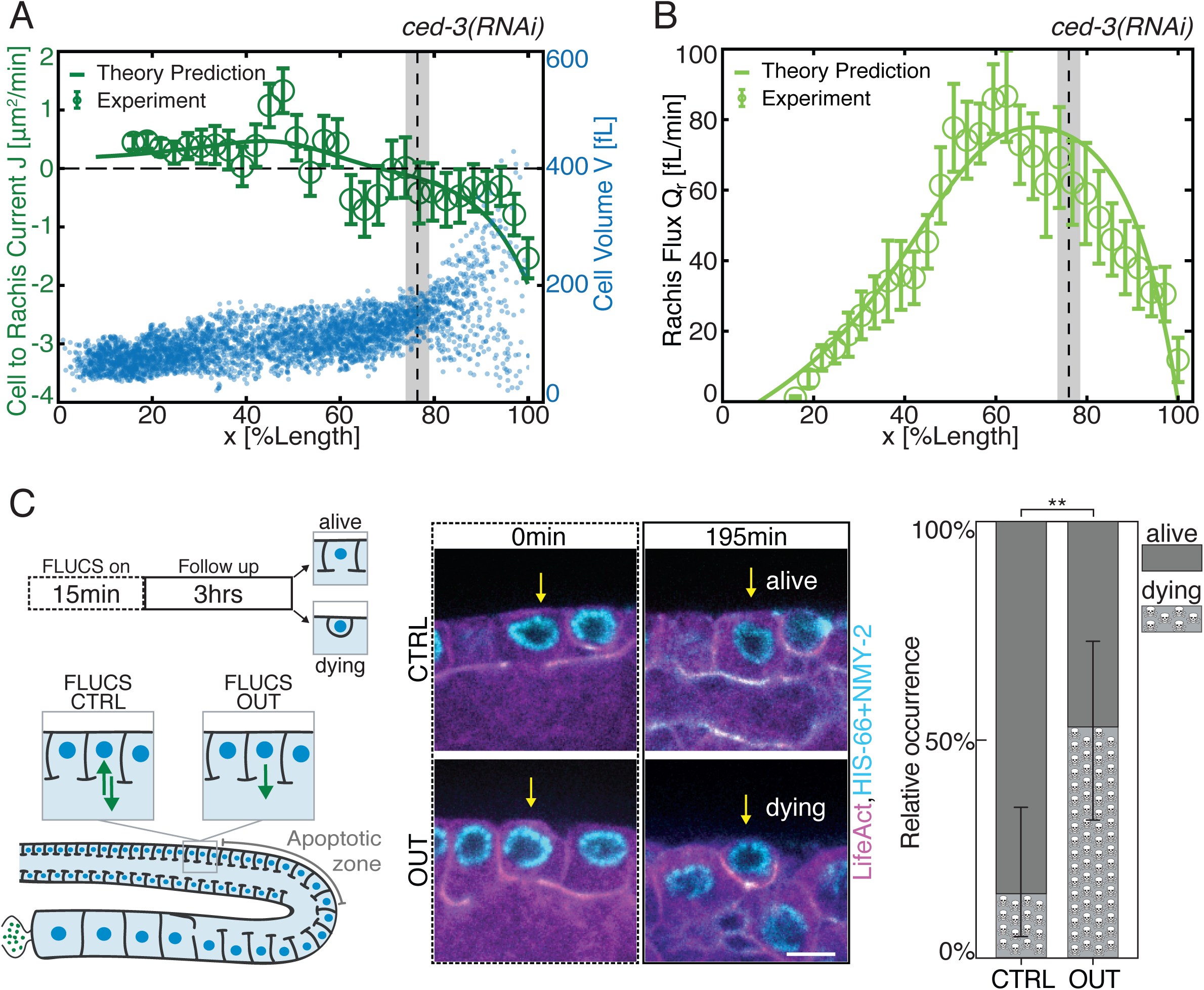
A hydraulic instability drives cell shrinkage which triggers cell death. **A**, Blue dots, cell volume *V* along the germline for 4030 germ cells from 7 gonad arms where apoptosis was inhibited by *ced-3(RNAi)*. Vertical dashed lines and grey bars in A and B denote the region of transition between germ cell growth modes for *ced-3(RNAi)* (Fig. 1B). Green open circles, germ cell to rachis current *J* in *ced-3(RNAi)* along the germline (see supplement). Green line, profile of the germ cell to rachis current *J* for *ced-3(RNAi)* as predicted from theory, using physical parameters and the profile of material uptake *S* obtained for the non-RNAi condition (see supplement, Figs. 1D, S2D). **B**, Rachis flux *Q*_*r*_ along the gonad for *ced-3(RNAi)* obtained from 9 gonad arms (green open circles, see supplement), together with the corresponding theory prediction (green line). **C**, FLUCS experiments. Left, schematic. Middle, representative fluorescence images before and after bidirectional FLUCS as a control (CTRL, top) and unidirectional FLUCS (OUT, bottom) (Magenta, Lifeact::mKate; cyan, GFP-NMY2 and Histone-Dendra). Right, relative occurrence of rachis bridge closure and rounding up within 3 hours after FLUCS CTRL or FLUCS OUT treatment; *\*\** denotes a p-value*<* 0.01. Scale bar; 5 *µm*. Error bars indicate the error of the mean at 95% confidence.

Our results suggest that the life and death decision in the gonad is a mechanical one. If so, it should be possible to bias the outcome of this decision via mechanical manipulation. In particular, artificially reducing the volume of individual germ cells should increase their likelihood to undergo apoptosis. We tested this prediction by unidirectional thermoviscous pumping (FLUCS, Focused-Light-indUced Cytoplasmic Streaming, (20)) for 15-20 minutes to pump germ cell cytoplasm out of individual germ cells through their rachis bridge, and monitoring the subsequent fate of the manipulated cells for three hours (Fig. 3C). As a control, we performed bidirectional FLUCS by rapidly switching between pumping cytoplasm into and out of individual germ cells, with an overall similar dosage of laser-light but without inducing net flow (Movie S5). In the control scenario, 14.3% of germ cells (3 out of 21) commence apoptosis within the following three hours, as judged by a characteristic rounding-up of apoptotic germ cells (5). This number compares favorably with rates of apoptosis in the unperturbed situation (Fig. S1C). In contrast, 52.6% of germ cells (10 out of 19) commence apoptosis within the three hours following unidirectional FLUCS (Fig. 3C, Movie S6). We conclude that a hydraulic manipulation to reduce the volume of individual germ cells results in an increased likelihood of commencing apoptosis. This lends credence to the statement that the life and death decision in the gonad is of mechanical nature.

Here we have shown that a hydraulic instability selects germ cells to either mature into oocytes or to donate their cytoplasm and die. Material uptake from the outside in the distal region leads to larger pressure in germ cells than in the rachis, which stabilizes germ cell volumes. This causes them to homogeneously grow while at the same time generating cytoplasmic flows through the rachis. Around 60% germline length the pressure difference between germ cell and rachis changes sign. This triggers a hydraulic instability that causes some germ cells to grow and others to shrink. Germ cell fate is then determined by size: shrinking cells undergo apoptosis while growing cells mature into oocytes. Note that the hydraulic instability amplifies small differences in germ cell volumes and redistributes material from smaller to larger cells. Hence, utilizing a hydraulic instability for the life and death decision in the gonad selects for larger and perhaps fitter cells while making use of the resources of the dying cells. The mechanism we have discovered here bases a cell fate decision on a hydraulic instability. This presents a robust alternative to biochemical switches usually invoked in cellular decision making processes (21-22).

## Acknowledgments

S.W.G. was supported by the DFG (SPP 1782, GSC 97, GR 3271/2, GR 3271/3, GR 3271/4) and the European Research Council (grant 742712). A.M. acknowledges support from the Joachim-Herz Stiftung. J.P. was supported by EMBO (ALTF 975-2019). B.T.L. acknowledges support through a National Science Foundation Graduate Research Fellowship. We thank the staff and students of the 2016 and 2017 MBL Physiology courses where this work was started, in particular J. Brzostowski, E. Eck, N. King, J. Lippincott-Schwartz, M. Morrison, C. Ott, B. Pennycook, R. Phillips, and N. Ratneyeke. We thank M. Mittasch for help with FLUCS, the light microscopy facilities of BIOTEC and MPI-CBG for support, and T.C. Middlekoop, L. Hubatsch, K. Ishihara for discussions and comments on the manuscript.

## Supplementary Materials

Materials and Methods

Figs. S1,S2

Supplemental Theory Notes

Movie Captions S1 to S6

